# Range Stability Predicts Lineage Persistence in a Morphologically Cryptic Ground Squirrel Species Complex

**DOI:** 10.1101/073320

**Authors:** Mark A. Phuong, Ke Bi, Craig Moritz

## Abstract

The processes responsible for patterns of cytonuclear discordance remain unclear. Here, we employ an exon capture dataset, demographic methods, and species distribution modeling to elucidate the impact of historical demography on patterns of genealogical concordance and discordance in ground squirrel lineages from the *Otospermophilus beecheyi* species complex. Previous studies in *O. beecheyi* revealed three morphologically cryptic and highly divergent mitochondrial DNA (mtDNA) lineages (named the Northern, Central, and Southern lineages based on geography) with only the Northern lineage exhibiting concordant divergence in nuclear markers. We show that these mtDNA lineages likely formed in allopatry during the Pleistocene, but responded differentially to climatic changes that occurred since the last interglacial (∼120,000 years ago). We find that the Northern lineage maintained a stable range throughout this period, correlating with genetic distinctiveness among all genetic markers and low migration rates between the other lineages. In contrast, our results support a scenario where the Southern lineage expanded from Baja California Sur during the Late Pleistocene and hybridized with the Central lineage, eventually driving the Central lineage to extinction. While high intraspecific gene flow among newly colonized populations eroded significant signals of Central ancestry from autosomal markers, male sex-biased dispersal in this system preserved signals of this past hybridization and introgression event in matrilineal-biased X-chromosome and mtDNA markers. Our results highlight the importance of range stability in maintaining the persistence of phylogeographic lineages, whereas unstable range dynamics can increase the tendency for lineages to interact and collapse upon secondary contact.

## Introduction

Understanding the spatial and temporal factors that influence the geographic distribution of genetic variation is a central goal in phylogeography (Avise 2000). One common phylogeographic pattern is cytonuclear discordance, or apparent differences in patterns of genetic differentiation among organelle DNA and nuclear markers (Toews and Brelsford 2012). Several non-mutually exclusive processes have been invoked to explain these phenomena, including simple stochastic variation in coalescent times across loci (Rosenberg 2003), local adaptation of distinct mitochondrial DNA (mtDNA) lineages (e.g., Ribeiro *et al*. 2011; Pavlova *et al*. 2013), and neutral introgression driven by range expansions during past climate oscillations (e.g., Cahill *et al*. 2013; Chavez *et al*. 2013). Despite the prevalence of cytonuclear discordance in natural systems, many previous studies have been unable to discern between several competing explanations for the described phylogeographic patterns (reviewed in Toews and Brelsford 2012).

Introgression during range expansions can produce patterns of cytonuclear discordance among closely related species through neutral demographic processes (Currat *et al*. 2008; Excoffier *et al*. 2009; Petit and Excoffier 2009), yet this mechanism has been difficult to test explicitly in empirical studies (but see Cahill *et al*. 2013). Range expansions occur through a series of founder events, ultimately creating clines of decreasing genetic diversity from the expansion origin to the expansion edge (Hewitt 1996; Slatkin and Excoffier 2012; Peter and Slatkin 2013). Range expansions increase the probability of genetic interactions between closely related species through range overlap (Currat *et al*. 2008); if interbreeding is possible when an expanding species collides with a local species, genes from the local species are expected to become massively introgressed into the invading species (Currat *et al*. 2008; Excoffier *et al*. 2009). Introgression of the invading genome with the local genome occurs in part because individuals from the colonizing species will initially exist at low densities, causing heterospecific matings to be more likely than conspecific matings (Currat *et al*. 2008). However, continued intraspecific gene flow within the invading species can erode this signal of introgression away, unless the mode of inheritance for particular genetic markers impedes or prevents this process (e.g., sex-linked markers, organelle genomes, Petit and Excoffier 2009; Cahill *et al*. 2013).

Although range expansions are ubiquitous in nature and may be responsible for patterns of cytonuclear discordance, testing of this demographic scenario has been hindered by a lack of (a) explicit analytical methods and (b) sufficient genomic data to properly test genetic predictions from range expansion theory. Few studies employ methods that explicitly use genetic variation and spatial information simultaneously to detect range expansions (but see Peter and Slatkin 2013; Jezkova *et al*. 2014; Pierce *et al*. 2014). Often, range expansions are inferred through other sources of information including fossil data (Hewitt 2004) or instances of introgression among closely related and morphologically distinct species (Wilson and Bernatchez 1998; Melo-Ferreira *et al*. 2005; Cahill *et al*. 2013). When range expansions are tested using genetic data, many studies employ methods that detect demographic expansions (e.g., Tajima’s D, Fu’s Fs, mismatch distributions) and qualitatively incorporate spatial information afterwards to infer colonization routes (e.g., Rowe *et al*. 2004; Perktas *et al*. 2011; Vences 2013) rather than explicitly incorporating geography into the genetic analysis. Further, range expansion theory makes explicit predictions about the relative degree of introgression among markers that experience differential levels of drift and intraspecific gene flow across the genome (Currat *et al*. 2008; Petit and Excoffier 2009). Accordingly, genome scale data is necessary to accurately infer these past demographic processes. Recently, Peter and Slatkin (2013) proposed a method based on a newly developed statistic ψ, the directionality index, to test for evidence of a range expansion and locate its origin. The method utilizes pairwise population comparisons of two dimensional site frequency spectra (2D-SFS, i.e., a summary of allele frequencies between two populations) to (a) detect changes in allele frequencies across geography caused by range expansions and (b) locate the expansion origin by explicitly incorporating spatial data within the same inference framework (Peter and Slatkin 2013). This new method, coupled with advances in sequencing technology that now allow the rapid obtainment of genomic data from populations, enables a robust examination of the impact of range expansions in generating patterns of cytonuclear discordance.

Here, we examine factors that may have led to cytonuclear concordance and discordance within the *Otospermophilus beecheyi* species complex. *O. beecheyi*, is a common and abundant ground squirrel in western North America that exhibits male-biased dispersal (Dobson 1979, 1982) and inhabits mesic habitats such as grasslands and agricultural areas (Howell 1938). Previous mtDNA sequencing of this group revealed three highly divergent, morphologically cryptic lineages within *O. beecheyi*, denoted as the Northern, Central, and Southern lineages based on geography (Álvarez-Castañeda and Cortés-Calva 2011; Phuong *et al*. 2014). These three lineages are parapatrically distributed, where the Northern lineage ranges from southern Washington to northern California, the Central lineage is found in the Sierra Nevada mountain region, and the Southern lineage extends from northern California to Baja California with an allopatric population in Baja California Sur (Fig. 1a, Phuong *et al*. 2014). These mtDNA lineages differ by 7-8% in sequence divergence, with the Northern and Southern lineages being more closely related to each other than either is to the Central lineage (Álvarez-Castañeda and Cortés-Calva 2011). In contrast, genetic analyses from a small set of nuclear microsatellites and sequenced loci supported the evolutionary independence of the Northern lineage, while the Central and Southern lineages were essentially genetically indistinguishable from each other (Phuong *et al*. 2014). Based on these previous results, the Northern lineage was elevated to the species, *Otospermophilus douglasii* (Phuong *et al*. 2014), but we refer to it here as the Northern lineage within the *O. beecheyi* species complex for consistency.

**Figure 1.**
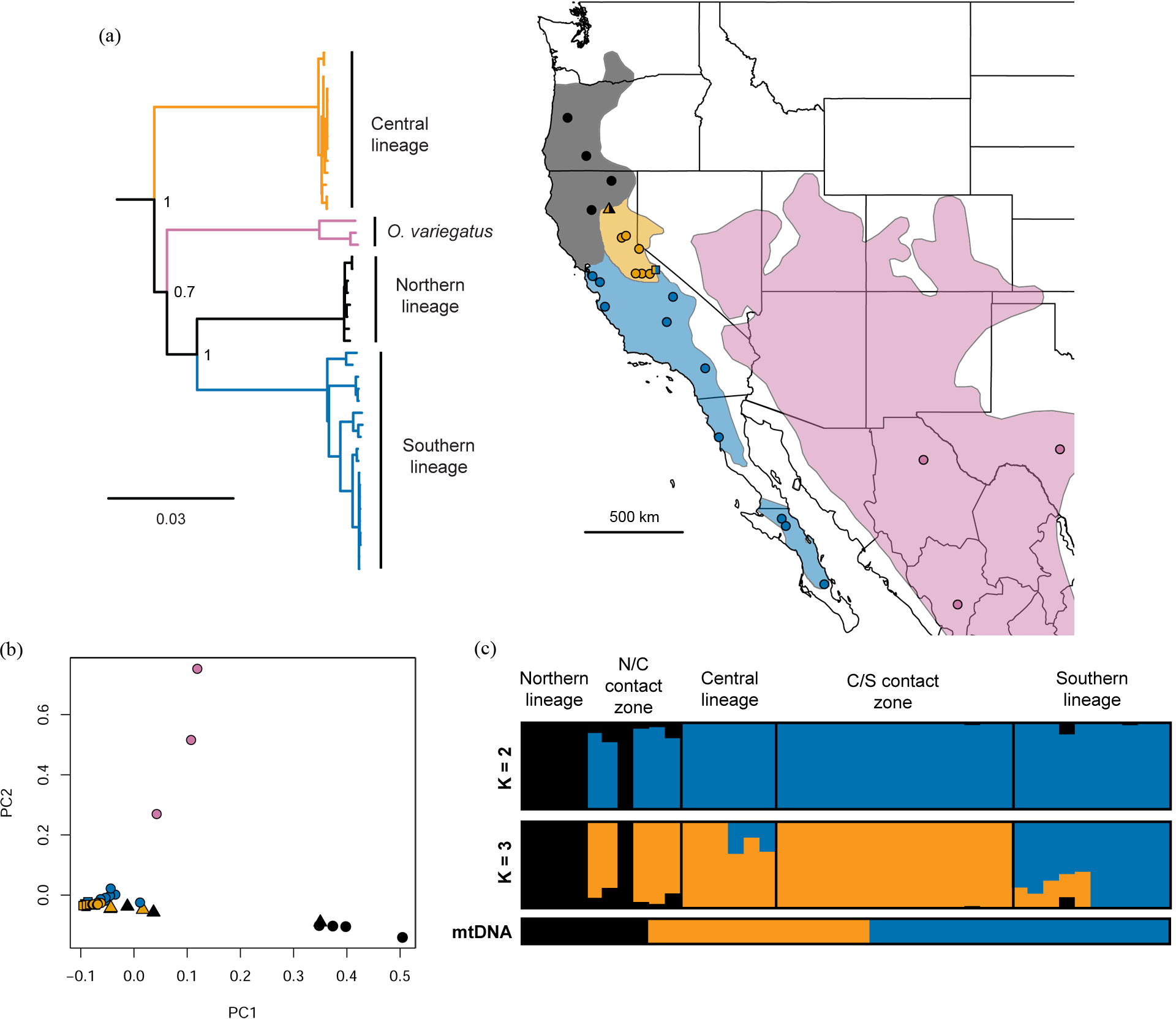
(a) Mitochondrial Bayesian phylogeny inferred from 13 protein coding genes and map of the western United States and Central America depicting the approximate geographic distribution of each lineage with colored shapes representing sampling localities. Labels at nodes represent posterior probabilities. The phylogeny is rooted with *Callospermophilus lateralis* (not shown), (b) Scatterplot of the first two principal components calculated from SNP data. Circles represent allopatric samples, triangles represent samples from the N/C contact zone, and squares represent samples from the C/S contact zone, as shown in the map in la. (c) ADMIXTURE population assignments at K=2 and K=3 from SNP data, with individuals (vertical bars) arranged by lineage and latitude. mtDNA lineage for each individual shown below the ADMIXTURE results.

To infer demographic processes that may explain why the Northern lineage exhibits concordance between mtDNA and nuclear markers while the Central and Southern lineages exhibit strong discordance, we employ an exon capture approach. We sequence several thousand genetic markers that vary in mode of inheritance (i.e., mtDNA, X-linked, autosomal) from individuals among the three lineages of the *O. beecheyi* species complex and for use as an outgroup, *O. variegatus* (a closely related species within *Otosopermophilus*). Then, we employ population genetic and demographic methods in conjunction with species distribution modeling to untangle the demographic history of this group. We also examine the role of selection on the mtDNA as an alternative explanation to patterns of cytonuclear discordance in *O. beecheyi*.

## Methods

### Genetic sampling & data collection

We obtained tissue samples from 44 individuals (i.e., eight Northern lineage, 14 Central lineage, 19 Southern lineage, three *O. variegatus*) from several institutions (Fig. 1a, Table S1). Six individuals were sampled from the contact zone between the Northern and Central lineages (N/C contact zone) and 15 individuals were sampled from the contact zone between the Central and Southern lineages (C/S contact zone, Fig. 1a). In addition, we included one sample of *Callospermophilus lateralis* as a distant outgroup with which to polarize ancestry of single nucleotide polymorphisms (SNPs, Table S1). We extracted genomic DNA using Qiagen DNeasy Blood and Tissue kits and prepared index-specific libraries detailed in Meyer and Kircher (2010).

To identify genes for target capture, we performed reciprocal blasts via blastx and tblastn (BLAST+, Altschup *et al*. 1990) between the annotated Beldingi’s ground squirrel (*Urocitellus beldingi*) transcriptome (Accession ##, published in this study) and the annotated transcriptome of the Alpine Chipmunk (*Tamias alpinus*, Bi *et al*. 2012) and identified loci that were > 7% divergent between the two species to increase the probability of recovering polymorphic loci in *O. beecheyi*. The *U. beldingi* transcriptome was sequenced and annotated following RNAseq lab protocols and bioinformatic steps described in Bi *et al*. (2012). We chose these two species for probe design because of their availability and close phylogenetic affinity to *O. beecheyi* (Harrison *et al*. 2003). We ultimately targeted ∼1.3 megabases consisting of 2305 nuclear protein coding genes and 13 mitochondrial protein coding genes on an Agilent SureSelect custom 1M-feature microarray. We inferred exon-intron boundaries by comparing *U. beldingi* transcripts with the genome of the thirteen-lined ground squirrel (*Ictidomys tridecemlineatus*) using EXONERATE (Slater and Birney 2005), identifying at least 3294 exons that were ≥ 200 bp in length. 249 of these exons were putatively X-linked and had corresponding orthologs on the X-chromosomes of human, mouse, rat, and dog. On the array, we used sequences from the *U. beldingi* transcriptome as our target sequence because it is more closely related to species within *Otospermophilus*. Each probe was 60bp in length and, to minimize edge effects, we tiled probes every 1bp for the first and last 6 bp of each exon, while probes were tiled every 4bp for the rest of the locus. We masked probes lying in short repeat or low complexity regions using the program repeatMasker (Smit *et al*. 2015) and repeated all probes three times on the array.

We pooled 50 libraries (45 made for this study and 5 used in another study) and hybridized this pool on the same array. Before hybridization, pooled libraries were denatured in the presence of excess blocking oligos and an excess 1:1 mixture of Cot-1 DNA isolated from *Mus musculus* and *O. beecheyi*. We isolated *O. beecheyi* Cot-1 DNA using the protocol described in Trifonov *et al.* (2009). We verified enrichment success using qPCR analysis and Bioanalyzer traces and sequenced all individuals on a single Illumina HiSeq 2000 lane with 100bp paired-end reads.

### Data filtration & assembly

We trimmed adapters and low quality bases using Trimmomatic (Bolger *et al*. 2014), merged overlapping paired-end reads using FLASH (Magoč and Salzberg 2011), removed reads with significant homology to human and bacterial (*E. coli*) genomes using bowtie2 (Langmead and Salzberg 2012), and removed duplicate and low complexity reads using custom Perl scripts. For each individual, we generated raw assemblies with a range of five k-mer values from 21–61 in ABySS (Simpson *et al*. 2009) and merged assemblies across k-mers using cd-hit (Li and Godzik 2006) and cap3 (Huang and Madan 1999)

To generate a mtDNA protein reference for each lineage, we first annotated mitochondrial proteins for one individual using DOGMA (Dual Organellar GenoMe Annotator, Wyman *et al*. 2004). Then, we chose one individual from each of the other lineages and performed a reciprocal blast via blastn to create lineage specific mtDNA protein references. Finally, we reconstructed protein coding sequences for each individual by aligning reads to these references using Novoalign (http://novocraft.com) and calling the base at each site directly from read depth. We masked sites that had less than 10X coverage or were not homozygous (minor allele frequency > 20%).

To generate a nuclear gene reference for all samples, we merged all final assemblies from every *Otospermohphilus* sample using cd-hit and cap3 and performed a reciprocal blast using blastn to identify contigs that matched the targeted exons. To account for chimeric contigs, we performed a self-reciprocal blast with blastn and removed contigs that had > 90% identity to each other. To generate a SNP dataset from the sequenced nuclear exons, we aligned all reads to the nuclear reference using Novoalign for each sample and identified variant sites using SAMtools (Li *et al*. 2009). To remove potentially paralogous sequences, we used a perl script (SNPcleaner, Bi *et al*. 2013) to remove contigs out of Hardy-Weinberg Equilibrium (HWE) from samples within each lineage that were outside the contact zones. We did not perform HWE filtering for *O. variegatus* due to low sample sizes. We kept variant sites with a minimum depth of 3X in at least 40 individuals and generated genotype likelihoods for every site within an individual using ANGSD (Korneliussen *et al*. 2014). We masked sites with ‘Ns’ when genotype posteriors were below 0.95. The bioinformatic scripts used in this study can be found at https://github.com/CGRL-QB3-UCBerkeley/denovoTargetCapturePopGen.

To assess the success of the capture experiment, we calculated (a) percent targeted bases covered by at least one read, (b) percent of reads aligned to intended targets and (c) sequence coverage for each individual.

### mtDNA analysis

We aligned and concatenated 13 mtDNA protein coding genes for all 45 individuals using MUSCLE (Edgar 2004) and inferred a phylogeny using MrBayes (Huelsenbeck and Ronquist 2001) under the partitioning scheme and substitution models estimated by PartitionFinder (Lanfear *et al*. 2012). We rooted the tree with *C. lateralis* and we sampled every 10,000 generations over 10 million generations with a 4 million generation burn-in. Further, because selection can explain genealogical discordance between mitochondrial and nuclear genealogies (Ballard and Whitlock 2004; Toews and Brelsford 2012), we tested for evidence of selection on mitochondrial proteins by comparing two site models (M1a and M2a) as implemented in PAML (Yang 2007).

### Nuclear population structure analysis

We characterized nuclear genetic structure in two ways: (a) we performed a principal component analysis (PCA) using smartpca from EIGENSOFT v4.2 (http://www.hsph.harvard.edu/alkes-price/software/), and (b) we inferred population assignment in ADMIXTURE, which uses a likelihood approach to estimate ancestry (Alexander *et al*. 2009). We ran ADMIXTURE for K=2 and K=3 populations. For these analyses, we randomly sampled 1 SNP per gene because both methods assume that SNPs are unlinked.

### Species distribution modeling

To infer how glacial cycles impacted range dynamics, we modeled each lineage’s distribution across several climate eras. We chose to model each lineage separately because, under our “expansion-introgression” model, we assumed that the mtDNA lineages once represented independent evolutionary entities that may have responded differentially to climatic shifts. We partitioned occurrence records from VertNet (vertnet.org; accessed on 30 January 2014) based on the geographic distribution of each mtDNA lineage. We included additional records (Álvarez-Castañeda and Cortés-Calva 2011; Phuong *et al*. 2014) that were not yet available on VertNet and we removed records if coordinate uncertainties were >10 km or were not reported. To avoid spatial autocorrelation in sampling, we thinned the data by randomly selecting one occurrence record per occupied cell in the bioclimatic layers using a custom R script (R Development Core Team, 2014). The final dataset included 161 unique records for the Northern lineage, 52 for the Central lineage, and 327 for the Southern lineage. We obtained 2.5 arc-minute resolution climate layers from the WorldClim database (Hijmans *et al*. 2005) under conditions for the present and the Last Glacial Maximum (LGM, ∼22,000 years ago) reconstructed under the Community Climate System Model. For the Last InterGlacial (LIG, ∼120,000 – 140,000 years ago), we reduced the resolution of the LIG climate layers to 2.5 arc minutes in ARCMAP from the available 30 arc-second resolution layers on the WorldClim database.

We generated species distribution models using Maxent 3.3.1 (Phillips *et al*. 2006) and parameterized Maxent using 7 of the 19 BIOCLIM variables that were not highly correlated with each other (Pearson correlation coefficient |r| < 0.7, Table S2). We used occurrence records of each mtDNA lineage and occurrence records of closely related ground squirrel taxa whose ranges overlap with the focal lineages in this study as pseudo-absences (Table S3, Figure S1). Occurrence points from closely related taxa were filtered with the same criteria described above. We trained all models on current conditions and projected species distributions onto climates representing each time slice. For each model, we executed 10 replicates and generated a consensus distribution map by averaging across all replicates. We evaluated model performance for the predicted contemporary distribution of each lineage by generating cross-validated area under the curve (AUC) values.

### Demographic analyses

To estimate the timing of divergence and the degree of nuclear migration between the lineages in this study, we used the program ∂a∂i to fit a three population isolation-with-migration model to the nuclear SNP data (Gutenkunst *et al*. 2009). ∂a∂i uses a diffusion-based approach to fit an expected demographic model to frequency spectrum data (Gutenkunst *et al*. 2009). Based on previous species tree reconstructions inferred from nuclear sequence data (Phuong *et al*. 2014), we constructed a model where the Northern lineage diverges first from an ancestral population, and then the ancestral population splits into the Central and Southern lineages. After divergence, migration is allowed to occur between lineages and happens continuously throughout their history. As we are primarily interested in historical, rather than current, introgression, we only included individuals outside of the contact zones for this analysis, except for one Northern lineage individual at the N/C contact zone that clustered with allopatric Northern lineage samples. We removed SNPs that were either (1) nonsynonymous variants, (2) found on X-linked exons, (3) adjacent to each other, or (4) had less than 5 diploid genotypes confidently called per lineage because of assumptions made by the ∂a∂i algorithm (Gutenkunst *et al*. 2009). We projected down to 10 allele copies per lineage for model fitting and performed the analyses on the folded 3D-SFS. To estimate parameter uncertainties, we used a nonparametric bootstrapping approach, where we generated 100 equal sized datasets by sampling SNPs with replacement from genes in the dataset using a Python script and repeated the ∂a∂i analysis. To convert coalescent units to real-time units, we used the average mammalian substitution rate of 2.2E-9 per base per year (Kumar and Subramanian 2002) and assumed the generation time in *O. beecheyi* to be 1 year (Dobson 1982).

Because our distribution modeling predicted that the Southern lineage expanded its range, we implemented the method described in Peter and Slatkin (2013) in R (R Development Core Team, 2014) to test for range expansion and to locate its origin on combined samples from the Central and Southern lineages, again excluding individuals at the contact zones. We included only synonymous nuclear variants and polarized the site frequency spectra with *C. lateralis*. We also investigated the effect of removing specific portions of the samples from the analysis to examine the impact on the method’s inference. We executed this analysis on the following groupings of samples: (a) only Central, (b) only Southern, (c) only Southern, without samples from Baja California Sur, and (d) Central + Southern, without samples from Baja California Sur.

Signatures of introgression due to a recent range expansion should be most evident in markers that have reduced intraspecific gene flow, such as in mtDNA or the X-chromosomes in mammals (Petit and Excoffier 2009; Cahill *et al*. 2013). Although high intraspecific gene flow can rapidly erode away strong signals of introgression in the autosomes, the reduced levels of intraspecific gene flow experienced by the matrilineal-biased X-chromosomes should retain elevated signatures of past introgression between species (Cahill *et al*. 2013). One prediction from these range dynamics is that we should expect elevated divergence in the X-chromosome relative to autosomes when comparing samples close to the expansion origin vs. the expansion edge because individuals in the recently colonized area will contain X-chromosomes of mixed ancestry (Cahill *et al*. 2013). To test this expectation, we generated the ratio of X-chromosome divergence to autosome divergence (X_*div*_/A_*div*_) by calculating uncorrected pairwise distances from the SNP dataset for female samples from all *O. beecheyi* lineages using a python script. Then, we categorized the pairwise comparison as either between (a) a sample from the origin vs. a sample outside of the origin, (b) only samples located outside of the origin, or (c) a Northern sample vs. a sample from another lineage. We performed comparisons including Northern lineage individuals to determine baseline X_*div*_/A_*div*_ values when there is no assumed history of introgression between the two individuals being compared. Based on results from the range expansion analysis, samples from the origin included two females from Baja California Sur, while samples outside the origin consisted of all other females from the Southern and Central lineages, exclusive of individuals from the contact zones. Using R, we performed an analysis of variance (ANOVA) to test for differences between these categories and conducted a post-hoc Tukey HSD test to determine which categories were significantly different from each other.

## Results

### Sequence capture statistics

We sequenced an average of 6.7 million reads per sample (Table S1). We recovered all mtDNA protein coding genes, sequencing an average of 99.9% of the bases. For each sample, approximately 7% of the reads mapped to the mtDNA protein coding genes, resulting in an average depth of 2495.6X (Table S1). We found corresponding orthologs for all nuclear exons targeted, generating a reference that was roughly 2.5 megabases in length (inclusive of flanking introns). We recovered an average of 97% of the bases with 8.7% of the reads mapping to the targeted exons, generating an average coverage of 17.6X (Table S1). We found high variance in percent reads on target and coverage in our dataset, despite attempts at equimolar pooling before hybridization with the array (Table S1). 354 of the targeted 3294 exons were removed due to being out of HWE. After further data filtering and genotype calling, we discovered 14,117 nuclear SNPs across 2361 loci (exons + flanking introns) that we used for subsequent analyses.

### mtDNA analysis

We recovered a mtDNA phylogeny with four divergent lineages corresponding to *O. variegatus* and the Northern, Central, and Southern lineages within the *O. beecheyi* complex (Fig. 1a). The topology and support values are consistent with previous studies that inferred relationships using a single mtDNA locus (Fig. 1a, Álvarez-Castañeda and Cortés-Calva 2011; Phuong *et al*. 2014). In particular, this more extensive dataset confirms that the Central lineage is more divergent from the Northern and Southern lineages than they are from each other. However, even with sequence data from 13 mtDNA protein coding genes, we were unable to infer the earliest diverging mtDNA lineage within *Otospermophilus* (Bayesian posterior probability = 0.7, Fig. 1a).

Both the nearly neutral model (M1a) and positive selection model (M2a) had equal likelihoods (ℓ = −20668.38131), and the positive selection model did not identify any mtDNA sites under positive selection (posterior probability < 0.95, Table S4). Thus, at this scale, the data does not support divergent selection as an explanation of the observed cytonuclear discordance.

### Nuclear structure analysis

The first two PCs explained 31.3% of the variance in the nuclear genetic data (Fig. 1b, Fig. S2). The PCA plot revealed three clusters that correspond to *O. variegatus*, the Northern lineage, and a mixture of Central and Southern lineage individuals (Fig. 1b). ADMIXTURE results showed that the Northern lineage was distinct at both K = 2 and K = 3, with evidence of localized hybridization at the N/C contact zone, where all but one individual showed ancestry from a separate population (Fig. 1c). The Central and Southern lineages are indistinguishable from each other at K = 2 (Fig. 1c). At K = 3, the Central and Southern lineages showed evidence of mixing between some allopatrically distributed individuals. All individuals at the C/S contact zone clustered with the Central lineage. (Fig. 1c). These results are consistent with population structure previously inferred from a significantly smaller set of microsatellites and sequenced nuclear loci (Phuong *et al*. 2014).

### Species distribution modeling

We were able to acceptably model the contemporary distribution of each lineage under current climatic conditions, as indicated by the mean AUC values for the Northern (AUC = 0.86), Central (AUC = 0.90), and Southern (AUC = 0.82) lineages. Models for the Northern lineage predicted suitable habitat in areas that closely match its current distribution across all time periods, suggesting range stability over the last 120 thousand years (Fig. 2a). For the Central lineage, models predicted a decline in suitable habitat within the Sierra Nevada (where it is currently distributed) since the LIG (Fig. 2b). The LIG and LGM models for the Central lineages also predicted habitat suitability in areas where it is not currently found today and we interpret these results with caution (Fig. 2b). For the Southern lineage, the LIG model predicted a restricted distribution in Baja California and along the coast of California, with habitat suitability increasing across California towards the present (Fig. 2c). This pattern suggests a potential range expansion of the Southern lineage from Baja California northwards into its current distribution in California.

**Figure 2.**
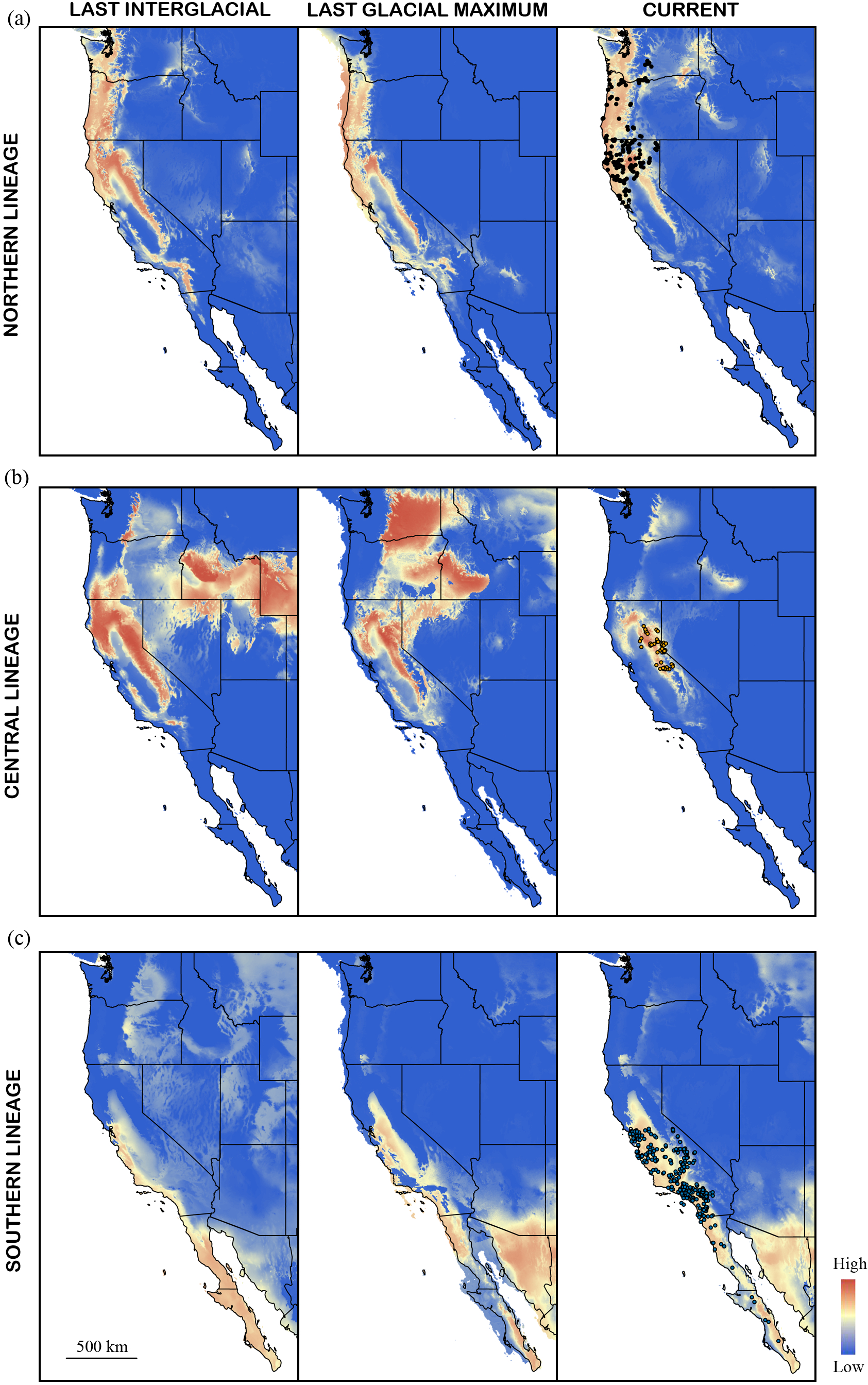
Species distribution models for the (a) Northern lineage, (b) Central lineage, and (c) Southern lineage from climates during the last interglacial (∼ 120-140 thousand years before present (kybp)), the last glacial maximum (∼20 kybp), and current conditions. Occurrence points used in distribution models are displayed on current climate maps and are colored black for the Northern lineage, orange for the Central lineage, and blue for the Southern lineage.

### Demographic analyses

Demographic modelling in ∂a∂i, conditioned on our phylogenetic model, revealed that the Northern lineage diverged from an ancestral Central and Southern lineage roughly 680,000 years ago, and the Central and Southern lineages diverged from each other approximately 180,000 years ago (Fig. 3, Table S5). The Southern lineage is inferred to have had the larger effective population size compared with the Northern and Central lineages, which had similar sizes to each other (Fig. 3, Table S5). Crucially, inferred migration rates between the Southern and Central lineages were an order of magnitude higher than between the Northern lineage and either the Central or Southern lineages (Fig. 3, Table S5). Although confidence intervals for the migration rates overlap (Table S5), the migration rate between the Southern and Central lineages was higher than migration rate estimates with the Northern lineage in every single bootstrap replicate (Fig. S3).

**Figure 3.**
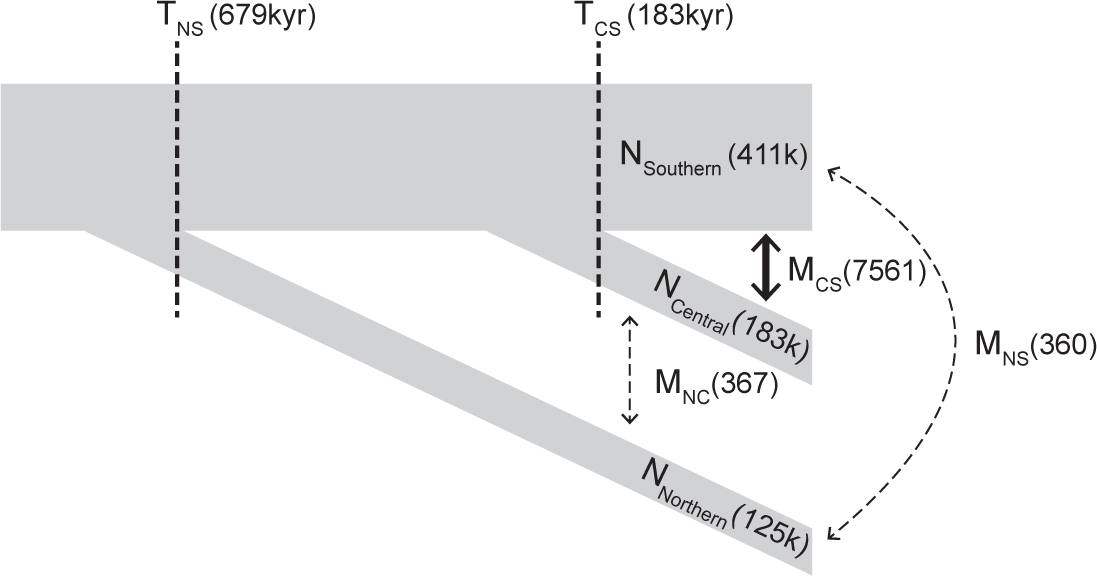
Schematic of the three population isolation-with-migration model with the Northern lineage diverging first and the Central and Southern lineage diverging thereafter. Point estimates reported in parentheses next to each demographic parameter: Divergence times T reported in thousands of years (kyr), effective population sizes N reported in thousands of individuals, and effective migration rates M reported in effective migrants per generation.

Using the directionality index, ψ, and the method developed by Peter and Slatkin (2013), we detected strong support for a range expansion (p < 0.0001) and located its origin to Baja California Sur (Fig. 4a) when including all Central and Southern lineage samples as a single population. This result is consistent with predictions from the modeling of potential lineage distributions through time. No other grouping of samples led to a significant signal for a range expansion (Fig. S4).

**Figure 4.**
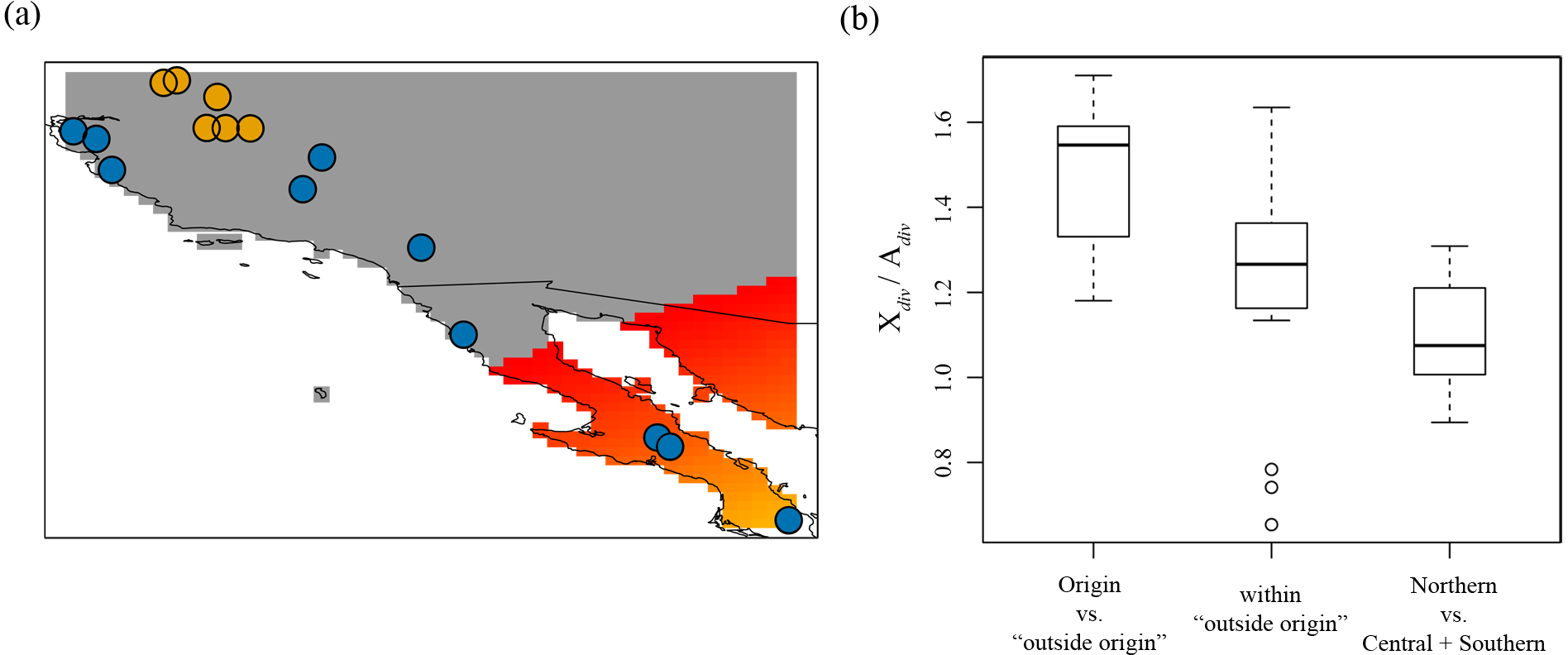
(a) Map of California and Baja California showing probability of origin of the range expansion. Grey represents no probability, and red represents high probability. Points on map represent samples used in the analysis, orange = Central lineage, blue = Southern linear (b) Ratio of X-chromosome divergence to autosome divergence when calculating genetic distances between (1) an individual from the origin vs. an individual outside of the origin (Origin vs. “outside origin”), (2) only individuals located outside of the origin (within “outsids origin”), or (3) a Northern individual vs. an individual from the Central or Southern lineage (Northern vs. Central + Southern).

The mean X_*div*_/A_*div*_ ratio for comparisons between samples from the expansion origin and expansion edge was 1.48 while comparisons from samples within the colonized area was 1.21 361 (Fig. 4b). The X_*div*_/A_*div*_ ratio when comparisons are made with a Northern lineage individual was 1.1 (Fig. 4b). We found significant differences in these ratios between the three categories (ANOVA, F = 16.37, p < 0.0001, Fig. 4b) and the Tukey HSD test indicated that X_*div*_/A_*div*_ was higher in the origin vs. “outside origin” category relative to the other two categories (p < 0.01).

## Discussion

Species histories are incredibly individualistic and complex; therefore, careful consideration of alternative explanations responsible for contemporary spatial patterns of genetic variation are critical to understanding the processes that have shaped the evolutionary history of species. First, we present our historical scenario to explain the results from our analysis of the *O. beecheyi* species complex. Then, we discuss potential weaknesses and inconsistences with our proposed model. In particular, we consider alternative processes that may have led to patterns within our dataset. Finally, we discuss the implications of our model in the context of broader evolutionary processes.

We propose that the mtDNA lineages within the *O. beecheyi* species complex represent evolutionary entities that were isolated from each other some time during the Pleistocene. Under this scenario, our demographic modelling placed these splits within the last several hundred thousand years (Fig. 3). In isolation, these lineages accumulated enough divergence to gain genetic distinctiveness in both mtDNA and nuclear DNA, as indicated by concordant genetic structuring in the nuclear genome for 3 out of the 4 mtDNA lineages within *Otospermophilus* (Fig. 1). Since the LIG, these lineages have responded differentially to climatic fluctuations which have shaped their present day genetic structure. Distribution models for the Northern lineage predicted a stable range since the LIG (Fig. 2a), which correlates with genetic distinction in both mtDNA and nuclear markers (Fig. 1) and a relatively lower rate of migration with the Central and Southern lineages (Fig. 3). These results suggest that the Northern lineage has maintained its evolutionary independence since isolation, despite demonstrated potential to hybridize with other lineages (as in the current contact zone, Fig. 1), because its distribution has been more stable across recent climatic shifts.

In contrast, Southern lineage distribution models suggested a recent range expansion from Baja California to its present day distribution in California (Fig. 2c), consistent with our genetic analyses showing statistical support for an expansion from the same locality (Fig. 4a). As the Southern lineage expanded across California, it invaded the range of the Central lineage and hybridized with it, causing Southern lineage individuals at the wave front to become massively introgressed by Central lineage genes (as predicted by theoretical simulations, Currat *et al*. 2008; Petit and Excoffier 2009). High intraspecific gene flow, as evidenced by relatively high migration rates among samples classified as Central and Southern individuals (Fig. 3), gradually eroded away significant signals of this hybridization event among the autosomes. Rapid autosomal erosion of Central lineage ancestry explains (a) why every Central and Southern individual – regardless of mtDNA haplotype – are genetically indistinguishable among autosomal markers (Fig. 1, Phuong *et al*. 2014) and (b) why the range expansion test was only significant when treating all Central and Southern lineage samples as a single evolutionary entity (Fig. 4a, Fig. S4). Due to male sex-biased dispersal in this system, erosion of Central ancestry on the X-chromosomes would proceed at a slower pace compared to autosomes while Central ancestry would never be erased from the maternally inherited mtDNA by mating events with male Southern colonizers. We find evidence for both these patterns, including (a) elevated levels of X-chromosome divergence between samples from the expansion origin vs. samples outside the origin (Fig. 4b) and (b) geographically structured and highly divergent Central lineage mtDNA (Fig. 1a). Over time, the Southern lineage likely drove the Central lineage to extinction because (a) theoretical simulations predict that the invading species will drive the local species to extinction if competition exists (Currat *et al*. 2008; Excoffier *et al*. 2009), (b) there is no strong contemporary signature of a genetically distinct Central lineage among autosomes(Fig. 1, Phuong *et al*. 2014), and (c) distribution models for the Central lineage suggest an apparent gradual decline in habitat suitability since the LIG (Fig. 2b). Further, replacement of the local species by the invading species is a common pattern across several empirical studies examining similar range dynamics among morphologically distinct taxa (e.g., Vachon 1998; Melo-Ferreira *et al*. 2005; Cahill *et al*. 2013) providing support for our scenario here among genetically distinct, but morphologically cryptic lineages.

Why did the Southern lineage invade the Central lineage, but not the Northern? We confirmed patterns from a previous study showing instances of localized hybridization and introgression at the N/C contact zone (Fig. 1, Phuong *et al*. 2014), suggesting that the Northern lineage is able to hybridize with other *O. beecheyi* lineages. The reasons explaining why hybridization with the Northern lineage is localized and not more widespread are unclear. Reproductive barriers, such as mating preference, hybrid inviability between heterospecific crosses, or ‘high-density blocking’ (Waters 2011), may limit the movement of Southern lineage individuals into Northern territory. Future studies examining reproductive isolation in this system may explain why the Southern lineage was not able to invade areas further north.

Several other scenarios could have occurred to generate the cytonuclear discordance pattern exhibited by the Central and Southern lineages. One alternative explanation is that because the nuclear genome evolves much slower than the mtDNA genome, not enough time has passed for these two lineages to gain distinction among nuclear markers. However, the lower nuclear genome substitution rate cannot explain why the Central mtDNA lineage, possibly one of the earliest diverging lineages in this complex, remains the only lineage in this complex to not have concomitant genetic distinctiveness among nuclear markers when all the other lineages do (Fig. 1). Indeed, one could argue that the data show slight signs of genetic distinctiveness between these two lineages, as evidenced by the ADMIXTURE K=3 showing some concordance between mtDNA and nuclear population assignment (Fig. 1c). However, range expansions cause geographic shifts in allele frequencies which can be detected by clustering algorithms (Peter and Slatkin 2013; Pierce *et al*. 2014), indicating that the apparent population distinction in the ADMIXTURE plot may be the result of the range expansion rather than the beginning stages of divergence. One other explanation is that selection on the mtDNA drove divergence among these lineages in the face of nuclear gene flow, as demonstrated in several phylogeographic studies of birds (e.g., Ribeiro *et al*. 2011; Pavlova *et al*. 2013). However, we did not find evidence for positive selection in our analyses (Table S4), suggesting that the mtDNA may better reflect historical demographic processes, rather than selective ones.

Here, we provide support for a rich demographic scenario that would otherwise go undetected if only autosomal markers were analyzed, adding to an extensive literature emphasizing the importance of matrilineal-biased markers in providing windows into ancient events in the past histories of organisms (Wilson and Bernatchez 1998; Melo-Ferreira *et al*. 2005; Currat *et al*. 2008; Cahill *et al*. 2013; Streicher *et al*. 2016). Specifically, we document the existence of a distinct Central lineage that likely no longer exists, but its past presence can only be detected because parts of the Central lineage genome (e.g., X-linked markers, mtDNA) have been ‘fossilized’ (Currat *et al*. 2008) or captured by the Southern lineage. While most studies that provide convincing evidence for a range expansion and introgression demographic scenario occur in systems where both lineages are still extant (Wilson and Bernatchez 1998; Melo-Ferreira *et al*. 2005; Cahill *et al*. 2013), there exists a small handful of cases where similar expansion-introgression events may have driven a once distinct evolutionary entity completely extinct (Bell *et al*. 2012; Singhal and Moritz 2012; Pereira *et al*. 2016). Although it is uncertain how often lineages become extinct and have portions of their genomes fossilized in invading colonizers through ‘expansion-introgression’ scenarios in natural systems, range expansions are thought to have occurred in the histories of most species (Excoffier *et al*. 2009). New demographic methods and sequencing techniques, such as the ones employed in this study, may lead to greater detection of this neutral demographic scenario in explaining patterns of cytonuclear discordance that may otherwise have been attributed to other factors such as selection (e.g., Toews and Brelsford 2012).

Our results highlight the importance of range stability in the maintenance of phylogeographic lineages whereas unstable range dynamics can lead to instances of lineage blending. As suggested in Singhal and Moritz (2013), one implication of this result is that climatically stable regions are therefore more likely to maintain and preserve the accumulation of lineages over time, whereas climatically unstable regions have a greater propensity for lineages to merge or become extinct. More broadly, these results align with ideas concerning the importance of climatic stability in the preservation of biological diversity at several scales of study, ranging from the accumulation of genetic diversity within populations in stable refugial areas (e.g., Hewitt 1996; Carnaval *et al*. 2009) to the concentration of species diversity in stable tropical climates (e.g., Mittelbach *et al*. 2007). Our study extends this general theme of stability maintaining diversity among genetically divergent and morphologically cryptic, phylogeographic lineages.

## Acknowledgements

For advice and discussions, we gratefully acknowledge, S Singhal, members of the KC Rowe laboratory, members of the Moritz laboratory, and members of the Ecology and Evolutionary Biology department at the University of California, Los Angeles. For samples, we thank the Burke Museum of Natural History and Culture, the Museum of Southwestern Biology, the Museum of Texas Tech University, the Grinnell Resurvey Project Team, the Deck family, ST Álvarez-Castañeda, TL Morelli, and JL Patton. We thank L Smith for laboratory advice, MCW Lim and DR Wait for assistance with the molecular work, and the Evolutionary Genetics Lab for providing a space to conduct the molecular work. We thank AB Smith for the R script to produce unique occurrence records. For the computational analyses, we thank the Alfaro laboratory for computing time. We thank RC Bell, MCW Lim, and S Singhal for insightful comments on earlier versions of this manuscript. This work was supported by the Museum of Vertebrate Zoology Undergraduate Biodiversity Award, a UC Berkeley Summer Undergraduate Research Fellowship, an NSF GRFP, an NSF GROW fellowship, a Chateaubriand fellowship, a Fulbright Fellowship, and an Edwin M. Pauley Fellowship to MAP, and a grant from the Moore Foundation to CM. This work used the Vincent J. Coates Genomics Sequencing Laboratory at UC Berkeley, supported by NIH S10 Instrumentation Grants S10RR029668 and S10RR027303.

---

**Figure S1.**
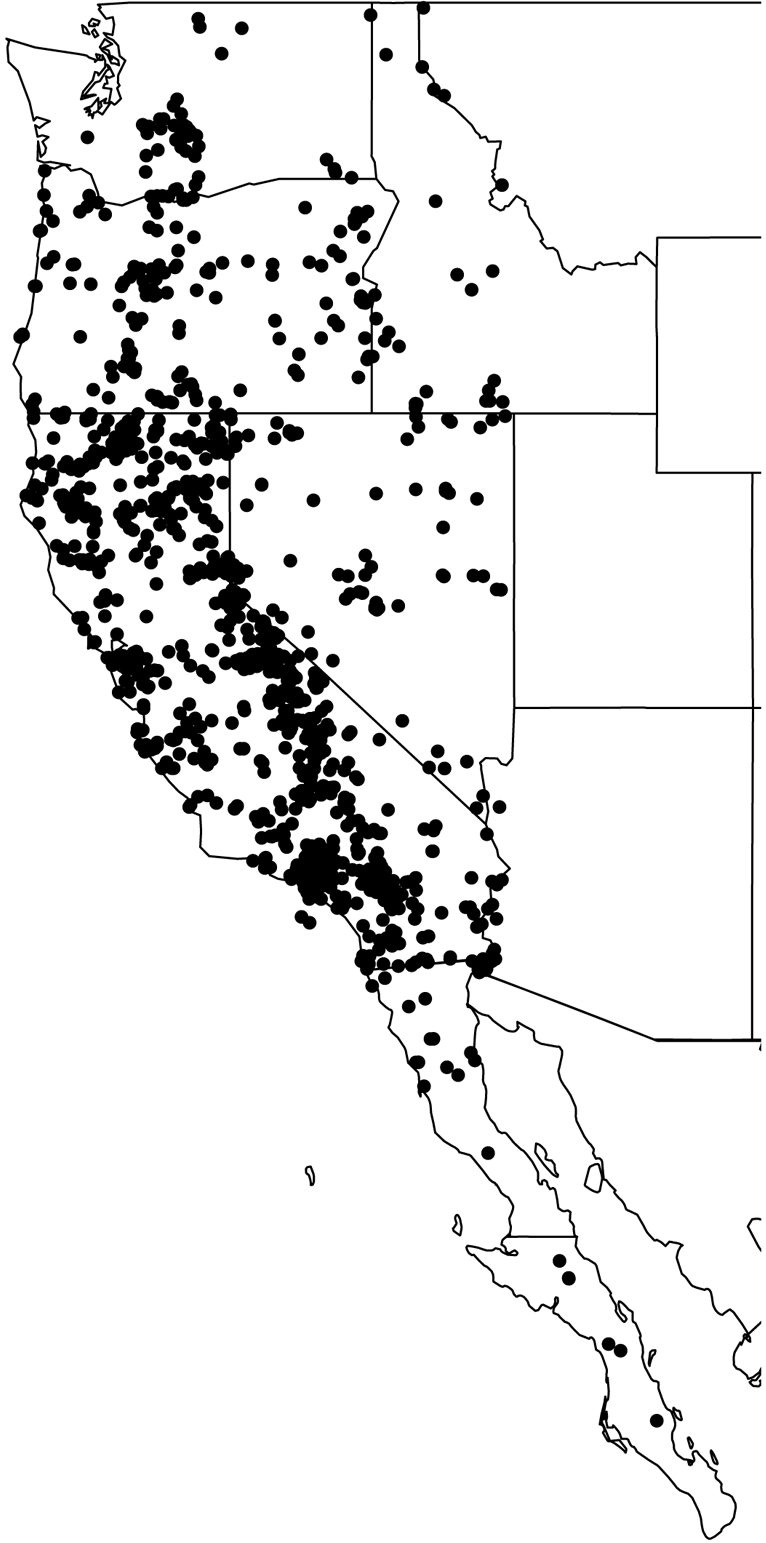
Pseudoabsences used in species distribution models.

**Figure S2.**
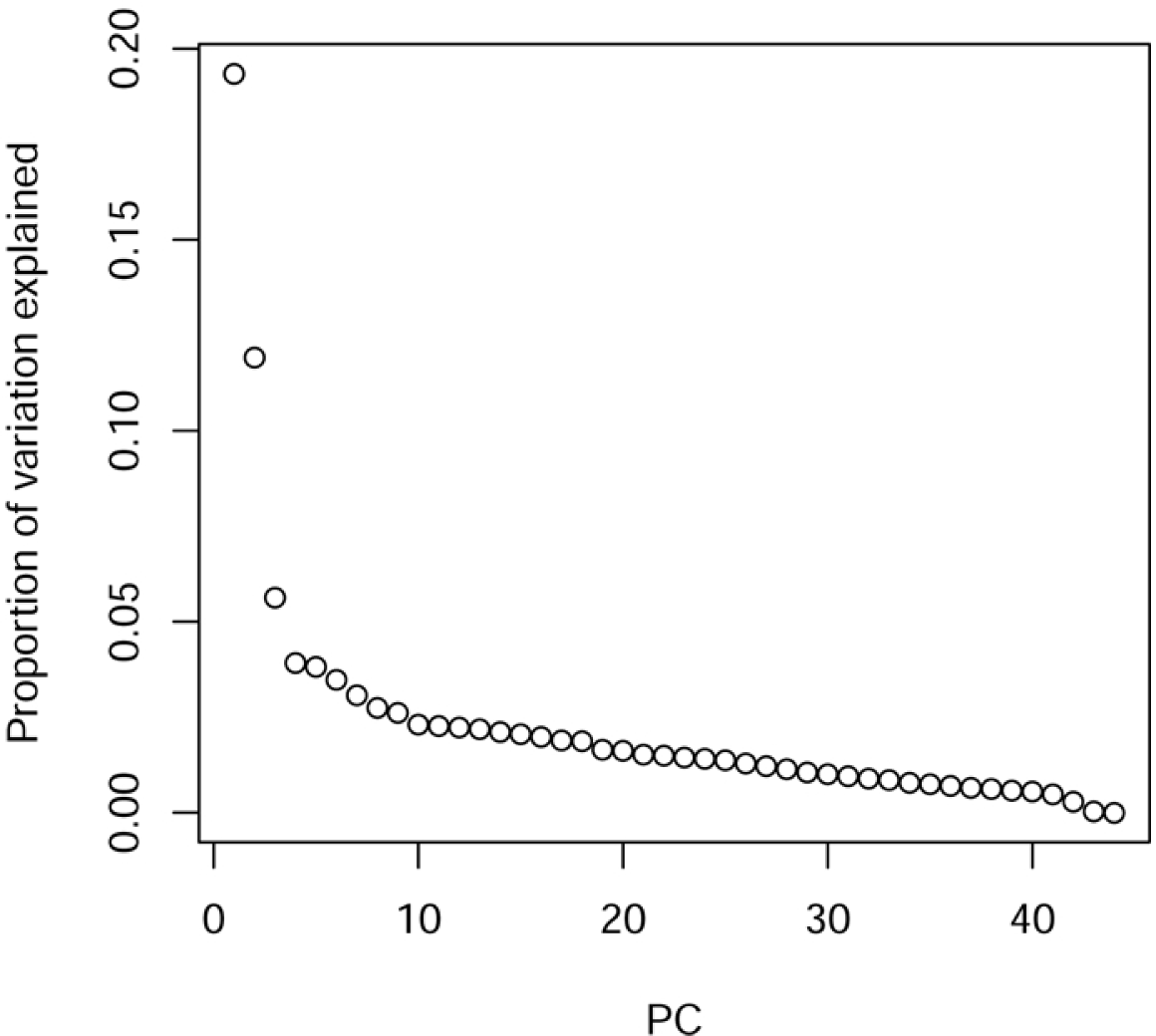
Proportion of variance explained by principal components generated from SNP data.

**Figure S3.**
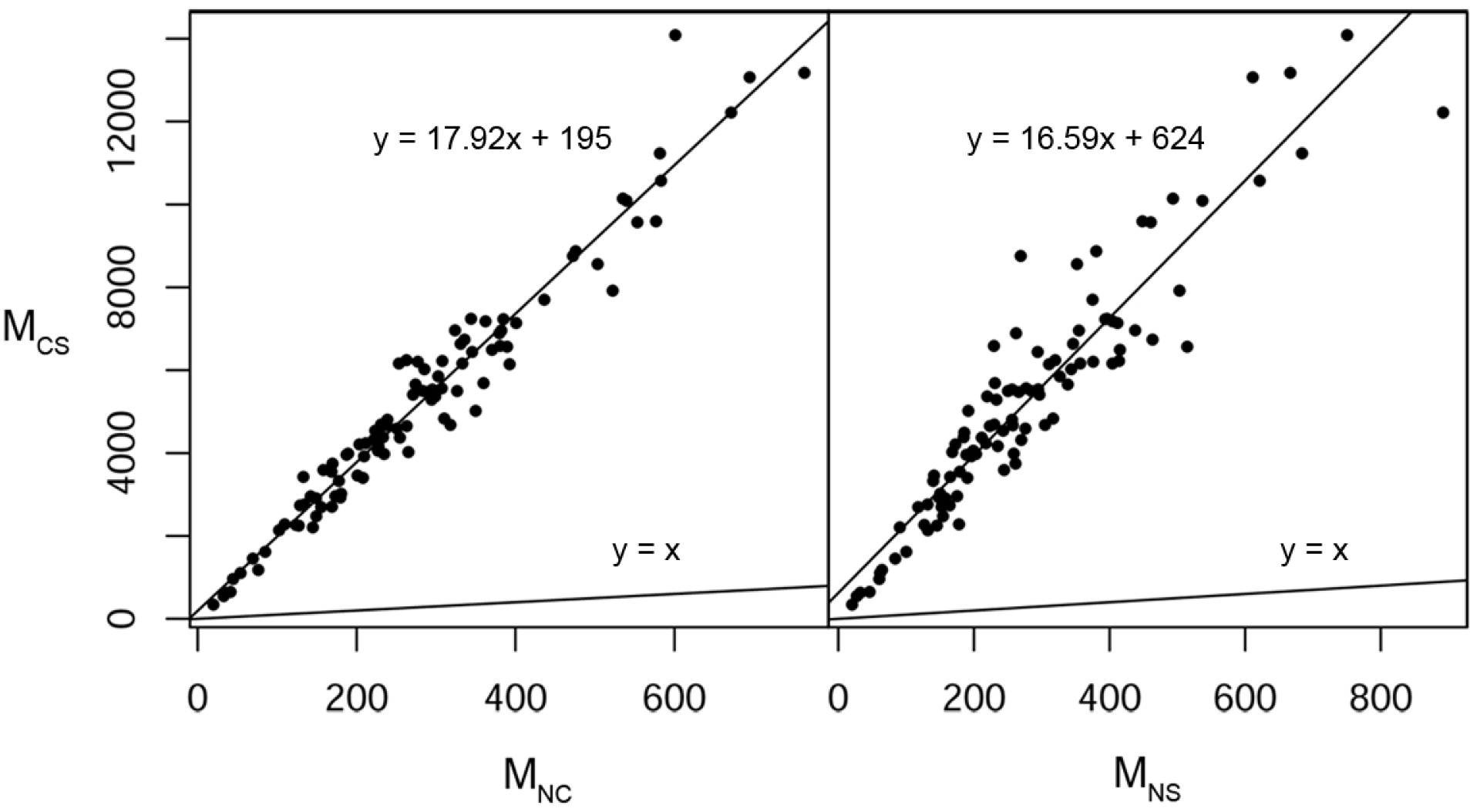
S3. Correlations between bootstrap estimated migration rates. Each point represents a bootstrap result, migration rates are reported as effective migrants per generation, and linear equations are shown. The y = x line is also shown.

**Figure S4.**
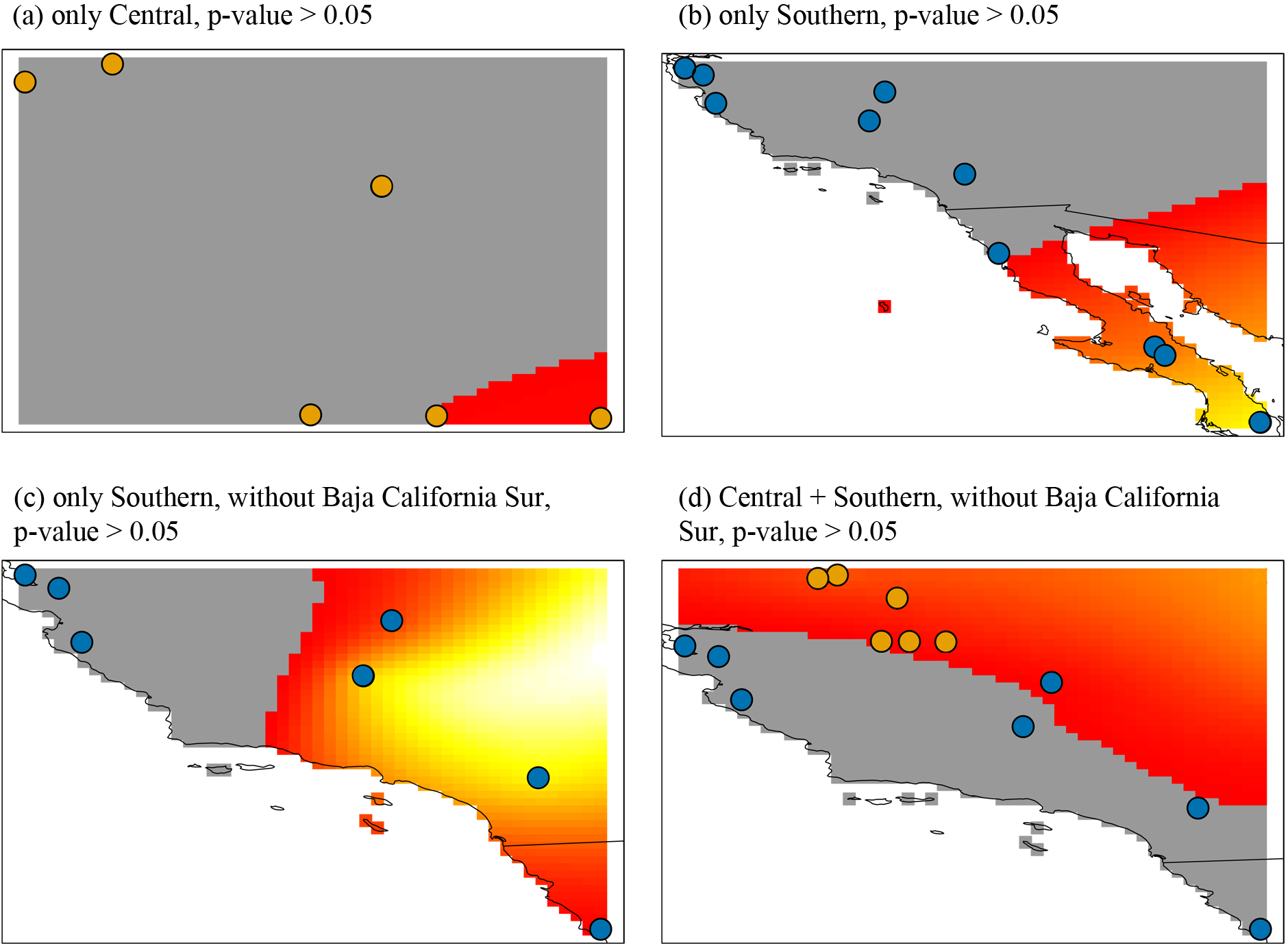
Range expansion tests for the following grouping of samples: (a) only the Central lineage, (b) Only the Southern lineage, (c) only the Southern lineage, without Baja California Sur, and (d) Central + Southern, without samples from Baja California Sur. Color on map represents probability of expansion origin, with grey representing low probability to white representing high probability. Samples are colored by lineage (orange = Central lineage, blue = Southern lineage), p-values for significant detection of a range expansion event are indicated.

**Table S1.** Samples used in this study and exon capture statistics. BM = Burke Museum; CIB = Centro de Investigaciones Biológicas del Noroeste; MVZ = Museum of Vertebrate Zoology; MSB = Museum of Southwestern Biology; TTU = Texas Tech University.

| Species | Institution | Sample ID | Latitude | Longitude | MtClade | SEX | # of reads | mtDNA |  |  | Nuclear exons |  |  |
| --- | --- | --- | --- | --- | --- | --- | --- | --- | --- | --- | --- | --- | --- |
|  |  |  |  |  |  |  |  | %bases | % on target | coverage | %bases | % on target | coverage |
| *C. lateralis* | MVZ | 207179 | 37.84631 | -119.63327 | *C. lateralis* | female | 8,489,208 | 100.0% | 9.7% | 3607.5 | 94.8% | 4.5% | 13.4 |
| *O. beecheyi* | MVZ | 201345 | 37.75139 | -119.79149 | Central | male | 6,505,546 | 100.0% | 7.1% | 2009.1 | 98.6% | 9.2% | 20.2 |
| *O. beecheyi* | MVZ | 207154 | 37.75606 | -120.08596 | Central | female | 5,050,286 | 100.0% | 6.8% | 1465.6 | 98.0% | 9.3% | 15.7 |
| *O. beecheyi* | MVZ | 207161 | 37.7409 | -119.40804 | Central | male | 4,523,914 | 100.0% | 5.0% | 949.3 | 97.3% | 9.4% | 14.0 |
| *O. beecheyi* | MVZ | 216232 | 37.89688 | -119.1298 | Central | female | 6,243,528 | 100.0% | 5.6% | 1488.4 | 98.6% | 9.5% | 19.3 |
| *O. beecheyi* | MVZ | 220666 | 40.40637 | -121.36086 | Central | female | 8,756,122 | 100.0% | 4.4% | 1567.9 | 99.4% | 11.6% | 32.1 |
| *O. beecheyi* | MVZ | 220667 | 40.40637 | -121.36086 | Central | female | 7,019,874 | 100.0% | 4.3% | 1257.0 | 98.8% | 10.0% | 22.6 |
| *O. beecheyi* | MVZ | 221757 | 39.23901 | -120.75351 | Central | male | 3,434,088 | 100.0% | 1.9% | 273.1 | 97.2% | 11.7% | 13.2 |
| *O. beecheyi* | MVZ | 224620 | 39.319381 | -120.549161 | Central | female | 4,043,082 | 100.0% | 6.6% | 1111.6 | 96.7% | 9.2% | 12.4 |
| *O. beecheyi* | MVZ | 226263 | 37.90028 | -119.12977 | Central | female | 3,272,870 | 100.0% | 6.7% | 909.0 | 96.1% | 10.0% | 11.0 |
| *O. beecheyi* | MVZ | 226264 | 37.90028 | -119.12977 | Central | male | 2,522,890 | 100.0% | 3.9% | 392.6 | 95.1% | 12.2% | 10.0 |
| *O. beecheyi* | MVZ | TLM101 | 38.77439 | -119.92002 | Central | male | 1,832,310 | 99.9% | 1.7% | 119.7 | 91.1% | 11.3% | 6.7 |
| *O. beecheyi* | MVZ | TLM26 | 37.98966 | -119.14156 | Central | female | 8,367,448 | 100.0% | 5.4% | 1868.0 | 99.2% | 10.2% | 27.4 |
| *O. beecheyi* | MVZ | TLM34 | 37.98966 | -119.14156 | Central | male | 7,311,450 | 100.0% | 1.2% | 341.5 | 99.2% | 12.9% | 29.7 |
| *O. beecheyi* | MVZ | TLM62 | 38.034836 | -119.213974 | Central | male | 4,961,404 | 97.5% | 0.3% | 65.4 | 98.2% | 11.2% | 17.4 |
| *O. beecheyi* | CIB | 12911 | 27.59 | -113.092 | Southern | female | 4,961,658 | 100.0% | 7.3% | 1563.6 | 97.0% | 7.4% | 12.5 |
| *O. beecheyi* | CIB | 12918 | 27.287 | -112.896 | Southern | female | 2,021,068 | 99.5% | 2.8% | 227.6 | 93.2% | 11.3% | 7.5 |
| *O. beecheyi* | CIB | 12928 | 24.871 | -111.055 | Southern | male | 4,960,810 | 99.9% | 5.1% | 1002.0 | 97.1% | 8.0% | 13.0 |
| *O. beecheyi* | MSB | 43115 | 30.966667 | -116.1 | Southern | female | 4,483,306 | 100.0% | 8.2% | 1601.1 | 97.0% | 8.3% | 12.7 |
| *O. beecheyi* | MSB | 43184 | 33.8238386 | -116.7563106 | Southern | female | 5,357,828 | 100.0% | 6.2% | 1377.9 | 98.1% | 9.8% | 16.7 |
| *O. beecheyi* | MVZ | 208499 | 37.90028 | -119.12977 | Southern | female | 3,930,888 | 100.0% | 6.0% | 987.3 | 95.5% | 7.4% | 9.9 |
| *O. beecheyi* | MVZ | 208500 | 37.95421 | -119.22578 | Southern | male | 18,250,922 | 100.0% | 14.7% | 12221.6 | 98.8% | 3.4% | 22.6 |
| *O. beecheyi* | MVZ | 216227 | 37.90028 | -119.12977 | Southern | female | 6,509,252 | 100.0% | 4.1% | 1093.4 | 98.7% | 9.5% | 20.1 |
| *O. beecheyi* | MVZ | 218030 | 37.6466861 | -122.1534062 | Southern | male | 6,277,164 | 100.0% | 3.1% | 784.6 | 98.4% | 9.5% | 19.9 |
| *O. beecheyi* | MVZ | 218700 | 37.396947 | -121.798483 | Southern | male | 9,533,900 | 100.0% | 13.4% | 5557.9 | 96.8% | 3.7% | 12.4 |
| *O. beecheyi* | MVZ | 220337 | 36.37851365 | -121.5568207 | Southern | male | 4,638,194 | 100.0% | 13.8% | 2680.6 | 87.0% | 3.0% | 5.8 |
| *O. beecheyi* | MVZ | 224848 | 36.78698 | -118.29751 | Southern | female | 3,716,938 | 100.0% | 7.2% | 1105.3 | 94.6% | 7.0% | 9.2 |
| *O. beecheyi* | MVZ | 224997 | 35.74516 | -118.59564 | Southern | female | 14,357,384 | 100.0% | 5.5% | 3287.4 | 97.6% | 3.1% | 14.3 |
| *O. beecheyi* | MVZ | 227066 | 37.90028 | -119.12977 | Southern | male | 7,577,124 | 100.0% | 8.8% | 2747.9 | 98.5% | 7.7% | 19.0 |
| *O. beecheyi* | MVZ | TLM11 | 38.01578 | -119.14993 | Southern | male | 4,577,268 | 99.7% | 0.8% | 141.8 | 98.6% | 13.1% | 19.4 |
| *O. beecheyi* | MVZ | TLM13 | 38.01578 | -119.14993 | Southern | female | 6,313,516 | 100.0% | 5.4% | 1322.1 | 98.5% | 9.1% | 20.1 |
| *O. beecheyi* | MVZ | TLM14 | 38.01578 | -119.14993 | Southern | female | 10,101,138 | 99.3% | 0.5% | 194.2 | 99.6% | 15.0% | 47.0 |
| *O. beecheyi* | MVZ | TLM179 | 38.04044 | -119.16781 | Southern | male | 4,550,400 | 99.4% | 0.8% | 152.4 | 98.2% | 12.0% | 17.5 |
|  |  |  |  |  |  |  |  | %bases | % on target | coverage | %bases | % on target | coverage |
| *O. beecheyi* | MVZ | TLM33 | 37.98966 | -119.14156 | Southern | female | 8,015,632 | 99.8% | 0.8% | 261.0 | 99.4% | 13.0% | 32.3 |
| *O. douglasii* | BM | 80593 | 42.62801 | -122.4496 | Northern | female | 9,392,804 | 100.0% | 10.1% | 3997.7 | 99.1% | 8.7% | 26.7 |
| *O. douglasii* | MVZ | 218052 | 40.3437 | -121.44671 | Northern | male | 4,518,458 | 100.0% | 12.0% | 2399.0 | 95.2% | 6.6% | 10.5 |
| *O. douglasii* | MVZ | 218054 | 40.34453 | -121.43214 | Northern | female | 22,741,600 | 100.0% | 23.2% | 25273.7 | 99.2% | 3.9% | 32.3 |
| *O. douglasii* | MVZ | 219576 | 40.38748 | -122.19799 | Northern | female | 9,699,344 | 100.0% | 16.3% | 7041.2 | 98.6% | 6.0% | 20.4 |
| *O. douglasii* | MVZ | 220665 | 40.44631 | -121.409 | Northern | female | 3,518,686 | 100.0% | 11.6% | 1783.1 | 94.3% | 7.4% | 9.0 |
| *O. douglasii* | MVZ | 223031 | 41.597655 | -121.244624 | Northern | female | 5,322,298 | 100.0% | 7.2% | 1625.9 | 97.5% | 9.1% | 15.6 |
| *O. douglasii* | MVZ | PLC1982 | 44.2112 | -123.3479 | Northern | male | 5,287,016 | 100.0% | 13.2% | 3174.2 | 96.7% | 6.9% | 12.9 |
| *O. douglasii* | MVZ | TLM45 | 40.28591 | -121.26085 | Northern | female | 4,848,144 | 100.0% | 1.2% | 243.4 | 98.6% | 12.8% | 19.6 |
| *O. variegatus* | TTU | 75454 | 24.03333 | -104.6667 | *O. variegatus* | NA | 2,578,774 | 99.9% | 8.9% | 966.9 | 91.0% | 6.6% | 6.3 |
| *O. variegatus* | TTU | 102298 | 30.46667 | -99.76667 | *O. variegatus* | NA | 11,093,746 | 100.0% | 7.3% | 3282.5 | 99.3% | 8.7% | 31.1 |
| *O. variegatus* | MSB | 199600 | 30.02755 | -106.30204 | *O. variegatus* | male | 8,232,884 | 100.0% | 17.9% | 6778.4 | 95.7% | 3.6% | 10.8 |

**Table S2.** WorldClim variables used in this study.

| Bioclim ID | Bioclim Variable |
| --- | --- |
| BIO1 | Annual Mean Temperature |
| BIO4 | Temperature Seasonality (standard deviation * 100) |
| BIO8 | Mean Temperature of Wettest Quarter |
| BIO9 | Mean Temperature of Driest Quarter |
| BIO12 | Annual Precipitation |
| BIO15 | Precipitation Seasonality (Coefficient of Variation) |
| BIO18 | Precipitation of Warmest Quarter |

**Table S3.** Species used as pseudo-absences for modeling.

| Species |
| --- |
| *Callospermophilus lateralis* |
| *Urocitellus beldingi* |
| *Urocitellus canus* |
| *Urocitellus mollis* |
| *Urocitellus saturatus* |
| *Xerospermophilus mohavensis* |
| *Xerospermophilus tereticaudus* |

**Table S4.**
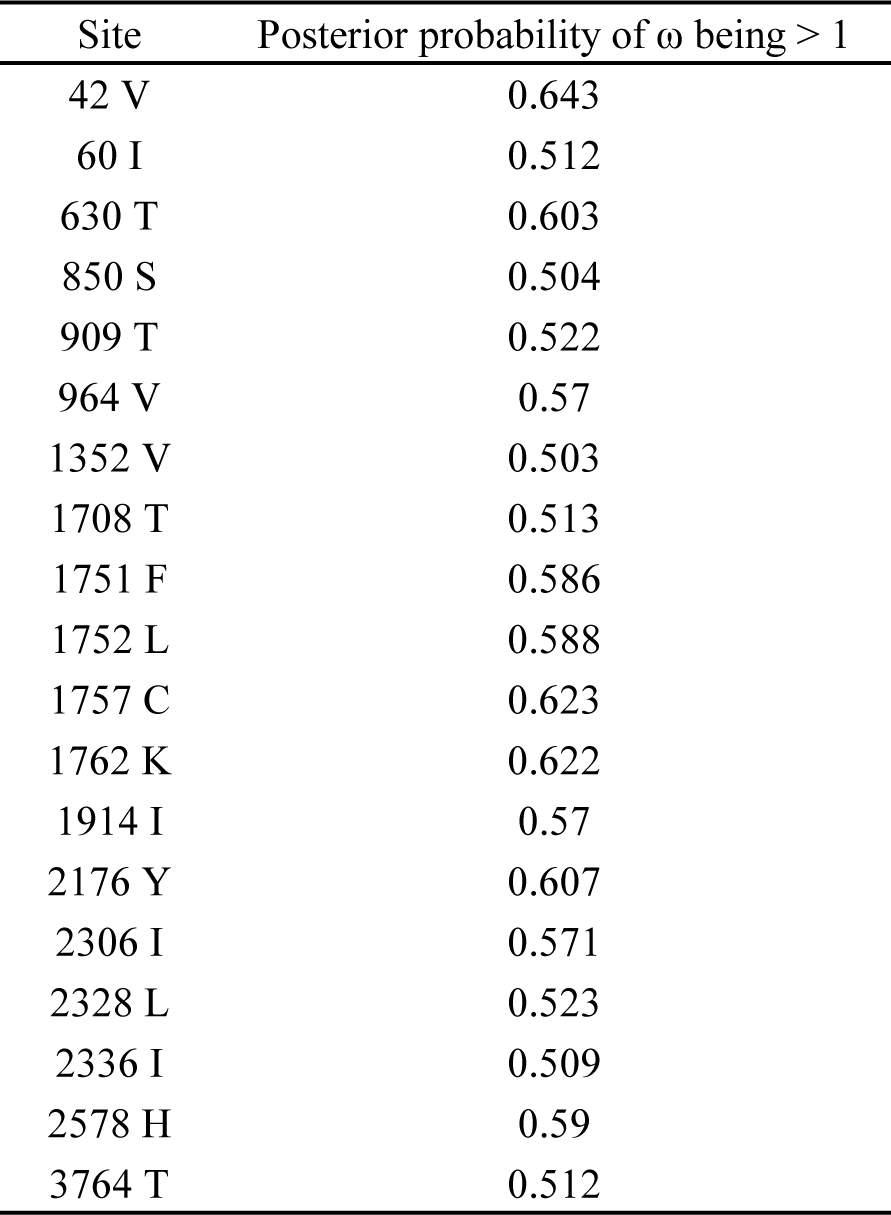
Sites under possible positive selection identified by PAML under the M2a model.

**Table S5.** Best-fit parameter estimates from dadi for the model schematic in Figure 3. Divergence time (T) reported in years, effective population size (N) reported in individuals, and migration rates (M) reported in effective migrants per generation. 95% confidence intervals, as calculated by # non-paramatic bootstraps, is reported in brackets.

| Parameter | Estimated value |
| --- | --- |
| $T_{NS}$ | 679265 [645368 - 722627] |
| $T_{CS}$ | 183190 [143829 - 214259] |
| $N_{Northern}$ | 125231 [115404 - 130796] |
| $N_{Central}$ | 173022 [151823 - 187761] |
| $N_{Southern}$ | 411308 [390570 - 455580] |
| $M_{NS}$ | 360 [0 - 588] |
| $M_{NC}$ | 367 [0 - 595] |
| $M_{CS}$ | 7561 [0 - 10859] |

